# Temperature alters specificity in a host-parasite interaction

**DOI:** 10.64898/2026.05.11.724370

**Authors:** Abbey Ramirez, Amanda Gibson

## Abstract

The Red Queen Hypothesis proposes that genetic variation is maintained in populations through antagonistic coevolution of hosts and parasites. A major assumption of the Red Queen Hypothesis is tight genetic specificity for infection. However, it has been argued that this genetic interaction of host and parasite (G_H_xG_P_) is sensitive to environmental context (G_H_xG_P_xE). Environmental change could accordingly disrupt coevolutionary oscillations on relevant time scales, calling into question antagonistic coevolution as a general and robust explanation for the maintenance of genetic diversity. To evaluate this critique, we used the plant-parasitic nematode *Meloidogyne arenaria* and its natural bacterial parasite *Pasteuria penetrans* to determine if specificity is altered by temperature. We exposed six isofemale host lines to five parasite sources at three ecologically relevant temperatures. We found that, at two of three temperatures, susceptibility to infection depended on the specific combination of host line and parasite source (G_H_xG_P_). This specificity varied across temperatures, consistent with a G_H_xG_P_xE effect. This three-way interaction was driven both by quantitative changes in the strength of specificity across temperatures and shifts in the susceptibility rankings of host-parasite combinations. Our study contributes a rare experimental test of a proposed challenge to the Red Queen Hypothesis and suggests the potential for environmental context to change host-parasite specificity.

## INTRODUCTION

The Red Queen Hypothesis proposes that host-parasite coevolution maintains genetic diversity through negative frequency-dependent selection. The key idea is that parasites adapt to specifically infect common host genotypes, conferring an advantage on rare host genotypes. This rare advantage prevents the loss of rare genotypes and limits the fixation of common ones (Haldane, 1949; Clarke, 1976; Jaenike, 1978; Bell, 1982). This long-standing hypothesis now has substantial support, arguing for an important role for coevolving parasites in maintaining genetic diversity and sexual reproduction (Lively, 1987; Chaboudez and Burdon, 1995; Lively and Dybdahl, 2000; Decaestecker *et al*., 2007; Wolinska and Spaak, 2009; Morran *et al*., 2011).

Despite the weight of evidence for the Red Queen Hypothesis, it has been challenged as a general explanation for the maintenance of diversity (Parker, 1994; Little, 2002; Otto and Nuismer, 2004). A key problem lies with the requirement of genetic specificity between host and parasite (G_H_xG_P_). Specifically, this genetic specificity may not be robust to environmental change, an inherent feature of natural settings. Red Queen models were founded on “Matching Alleles” specificity, which models self/nonself recognition: hosts kill parasite genotypes that are recognized as non-self, but fail to recognize and thus succumb to infection by parasite genotypes that mimic self (“a match”) (Hamilton, 1980, 1993; Frank, 1997; Grosberg and Hart, 2000; Agrawal and Lively, 2002; Lambrechts, Fellous and Koella, 2006; Dybdahl, Jenkins and Nuismer, 2014). Thus, parasites evolve to “match” the common host genotype at the expense of infecting rare host genotypes, driving negative frequency-dependent selection. Coevolutionary models have since established that Red Queen dynamics and the maintenance of genetic diversity do not require this strict one-to-one specificity and can occur under alternate infection matrices (e.g., gene-for-gene); however, some form of G_H_xG_P_ specificity is required (Agrawal and Lively, 2002; Engelstädter and Bonhoeffer, 2009; Kwiatkowski, Engelstädter and Vorburger, 2012; Engelstädter, 2015). Supporting this requirement, there is ample empirical evidence that infection success depends on the interaction of host and parasite genotype (Lively, 1989; Carius, Little and Ebert, 2001; Koch and Schmid-Hempel, 2012; Rouchet and Vorburger, 2012).

A remaining challenge is that negative frequency-dependent selection by parasites requires that this specificity be stable and persistent. Yet the genetic interaction between hosts and parasites may be sensitive to the environment (G_H_xG_P_xE) and thus liable to change on relevant temporal scales (Blanford *et al*., 2003; Thomas and Blanford, 2003; Lazzaro and Little, 2008; Laine, 2009; Wolinska and King, 2009). Environmental variation could disrupt host-parasite specificity in two ways. First, the relative performance of combinations of host and parasite genotypes could change across environmental contexts, such that host-parasite combinations with relatively high infection in one context have relatively low infection in another. This effect is referred to as a shift in rank order. These shifts in fitness rankings can weaken or eliminate rare advantage, if rare host genotypes are under-infected only in certain contexts. Second, the strength of specificity could vary across environmental contexts, with greater divergence among host-parasite combinations in some contexts than in others. This effect is expected to generate variation in the magnitude of the advantage of rare genotypes (Mostowy and Engelstädter, 2011). These two scenarios – rank order shifts or changes in the strength of specificity – are not mutually exclusive, and in either case, environmental variation could destabilize G_H_xG_P_ specificity and undermine negative frequency-dependent selection.

There have been few empirical tests for G_H_xG_P_xE interactions. Wolinska and King (2009) reviewed studies examining how environmental context influences host-parasite interactions and found that while 31 studies reported significant genotype-by-environment (i.e., G_H_xE or G_P_xE) effects (Mitchell *et al*., 2005; Laine, 2007), only one tested for a G_H_xG_P_xE interaction. This study, Tétard-Jones *et al*. (2007), paired six barley genotypes with four different aphid clones and measured barley height and aphid population growth. The specific interaction of barley and aphid genotypes for these traits was significantly altered by the addition of the rhizobacteria *Pseudomonas aeruginosa* (i.e., a G_H_xG_P_xE interaction). Since Wolinska and King’s (2009) review, a few additional studies have detected G_H_xG_P_xE interactions, including for the interaction of the chestnut blight fungus *Cryphonectria parasitica* with its hyperparasitic virus (Bryner and Rigling, 2011), the Pacific oyster *Crassostrea gigas* with the bacteria *Vibrio splendidus* (Wendling, Fabritzek and Wegner, 2017), and the bumblebee *Bombus terrestris* with its trypanosome parasite *Crithidia bombi* (Sadd, 2011). In contrast, no support was found in the highly specific interactions of the cladoceran *Daphnia magna* with its bacterial parasite *Pasteuria ramosa* (Vale and Little, 2009; Duneau *et al*., 2011), or the black bean aphid *Aphis fabae* and its bacterial endosymbiont *Hamiltonella defensa* with the parasitoid wasp *Lysiphlebus fabarum* (Cayetano and Vorburger, 2013; Gimmi and Vorburger, 2021). These conflicting results, combined with the low number of empirical tests, leave uncertain whether environmental disruption of specificity poses a legitimate challenge to the Red Queen Hypothesis.

We aimed to address this gap by testing for a G_H_xG_P_xE effect in the interaction of the root-knot nematode *Meloidogyne arenaria* and its bacterial parasite *Pasteuria penetrans*. Endospores of *P. penetrans* attach to the cuticle of *Meloidogyne* nematodes, then germinate, extending a germ tube into the body of the host and replicating vegetatively (Chen and Dickson, 1998). The attachment step is required for subsequent infection, and it depends upon the specific interaction of host and parasite genotype, such that the outcome of infection cannot be predicted by the main effects of host or parasite alone (i.e., a G_H_xG_P_ interaction) (Stirling, 1985; Channer and Gowen, 1992; Mundim and Gibson, 2022). *Pasteuria penetrans* is a close relative of *P. ramosa*, which shows strong genetic specificity to its host *D. magna* that is consistent with the Matching Alleles model (Luijckx *et al*., 2011, 2013; Bento *et al*., 2017). This solid foundation for G_H_xG_P_ specificity in the *P. penetrans-M. arenaria* interaction positioned us to detect a G_H_xG_P_xE interaction if present. We focus on the effect of temperature, as it is a prominent environmental variable that may strongly influence interactions between ectothermic hosts and their pathogens (Blanford *et al*., 2003).

In this study, we tested for a G_H_xG_P_xE interaction by measuring endospore attachment for 30 combinations of host and parasite genotypes in three temperatures that are ecologically relevant for *M. arenaria* and *P. penetrans* (Hatz and Dickson, 1992; Chen and Dickson, 1997; Dávila-Negrón and Dickson, 2013). Mean attachment rates of *P. penetrans* to *Meloidogyne* were previously shown to vary over these temperatures (Freitas, Mitchell and Dickson, 1997). We extended this work by assessing whether temperature affects the specificity of attachment. We anticipated one of two potential outcomes. The finding of a G_H_xG_P_xE interaction for attachment would indicate that host-parasite specificity was not stable across two or more temperatures, consistent with the idea that environmental context has the potential to undermine the rare advantage that is critical to the Red Queen Hypothesis. The finding of an overarching G_H_xG_P_ interaction, with no G_H_xG_P_xE interaction, would indicate that host-parasite specificity was constant across the three temperatures, indicating stable genetic specificity. In evaluating these two potential outcomes, we address an ongoing challenge to the Red Queen Hypothesis.

## METHODS

### Study System

Our host in this study was the plant parasitic nematode *Meloidogyne arenaria. Meloidogyne* is a genus of plant parasitic nematode that causes billions of dollars in crop damage annually (Chitwood, 2003; Abad *et al*., 2008). *Meloidogyne arenaria* is an asexual species that infects peanuts (*Arachis hypogaea*) and many other crop plants (Rich *et al*., 2009). These nematodes establish in the roots of plants and prevent the uptake of nutrients and water, causing stunted growth, low yield, and death. Infection is initiated by second stage juveniles (J2). These J2s hatch in the soil, then search for plant root tips. Once they enter a root, the nematodes establish feeding sites where they develop into bulbous, sedentary adult females. They then reproduce by depositing a single egg mass on the surface of the root. This process takes approximately 30-40 days, and each egg mass can contain up to 1000 eggs (Evangelina García and Sánchez-Puerta, 2012). Because *M. arenaria* reproduces via mitotic parthenogenesis, offspring in an egg mass are genetically identical to their mother and to one another (Chitwood, 2003; Blanc-Mathieu *et al*., 2017). This reproductive mode allows us to propagate lineages clonally by establishing isofemale lines from single egg masses collected in the field.

Our parasite in this study was *Pasteuria penetrans*, an endospore-forming bacterium that is an obligate parasite of *Meloidogyne* species, including *M. arenaria* (Bishop, 2011). Endospores of *P. penetrans* attach to the cuticle of J2 hosts as they migrate through the soil. Once the nematode enters the root, establishes a feeding site, and begins to mature, attached endospores send a germ tube through the cuticle and replicate vegetatively in the body of the host, filling it with endospores. Infected hosts make few to no offspring. New endospores enter the soil after the plant roots decompose, and the host cadaver breaks open.

In this study, we specifically focus on the attachment of endospores to the nematode cuticle. This step of the infection process is most relevant to evaluating a G_H_xG_P_xE interaction for two reasons. First, attachment is the first step of infection: it captures the initial interaction of the host and parasite and is required for subsequent development of the parasite. Second, attachment is considered the primary step at which specificity is expressed in the interaction of *M. arenaria* and *P. penetrans*. There is indeed evidence of strong specificity at this step (Stirling, 1985; Channer and Gowen, 1992; Mundim and Gibson, 2022), and it is thought to arise from variation in the binding proteins of the parasites and receptors on the host cuticle (Davies and Opperman, 2006; Davies, 2009). Moreover, in infections of *Daphnia magna* by the related pathogen *Pasteuria ramosa*, attachment is the infection step that depends most strongly on the interaction of host and parasite genotypes and is thus the most likely to drive coevolution (Duneau *et al*., 2011).

### Host lines

We propagated isofemale lines of *M. arenaria* on eggplants (*Solanum melongena,* cv. Black Beauty) in the greenhouse. Prior to inoculation with *M. arenaria* eggs, we started eggplant seedlings in incubators set to 28°C, then after ∼three weeks transplanted them into small pots filled with 80% sand and 20% topsoil. We reared seedlings for ∼two weeks in the greenhouse, then transplanted them into larger pots and allowed them to establish for two more weeks before inoculating them. Plants were kept at a 16:8 light cycle at 23.9–26.7°C during the day and 21.1–24.4°C during the night.

To establish our *M. arenaria* lines, we collected egg masses of *M. arenaria* from United States Department of Agriculture (USDA) peanut fields in Tifton, GA in August of 2022 and 2023. We collected peanut roots from two different farms ∼ 8km apart and from 1-5 fields per farm (Table A1). We later processed the roots at the University of Virginia. We washed roots and identified egg masses by dyeing the roots in 20% red food coloring for 15 minutes (Thies, Merrill and Corley, 2002). The red dye stained the egg masses, making them easy to locate. We inoculated the ∼8-week-old eggplants with a single egg mass and allowed the isofemale lines to establish and reproduce for at least four months. We collected approximately 120 isofemale lines over this sampling period. For this experiment, we used six lines that were each sampled from a unique field to maximize the probability that lines were genetically distinct.

### Parasite sources

We opted to conduct this study with “parasite sources” rather than isolates. *Pasteuria penetrans* cannot be cultured outside the host, so establishment of one isolate requires multiple generations of culturing on *Meloidogyne* hosts reared on plants in the greenhouse (Andras *et al*., 2020). Power to detect a three-way interaction requires substantial replication, including multiple parasite lines crossed with multiple host lines. To ensure we could achieve a sufficient level of replication for the parasite, we elected to instead collect parasite sources directly from the field. Soil samples collected from the field could contain more than one genetically distinct *P. penetrans* isolate, so we took measures in our sampling to limit the diversity within a source and increase the diversity between sources. This approach resembles related tests of specificity and local adaptation that use populations of parasites (Hufbauer and Via, 1999; Lively and Dybdahl, 2000; Prugnolle *et al*., 2006).

We collected *P. penetrans* endospores in August 2023. Endospores were collected from the same farms as the host lines, but not in the same locations (Table A1). For each source of *P. penetrans,* we took three soil cores from around the roots of one infected peanut plant, combined the three cores into a single mixed sample (i.e., a “source”), and then completely dried the soil to kill any nematodes present. With this approach, we aimed to sample a large number of endospores representing variants circulating naturally in the same region as our host lines. Spores of *P. penetrans* are non-motile, so we expect little dispersal in soil, with limited movement between fields over the course of a growing season. We accordingly collected one source per field to maximize the potential for genetic differentiation between sources. By limiting the sampling scale for each source, we aimed to minimize the diversity of *P. penetrans* isolates per source. Each parasite source may nonetheless contain multiple parasite genotypes and thus the parasite “genotype” in G_H_xG_P_xE should be thought of as parasite populations, rather than isolates.

Parasite sources likely differed in their abundance of endospores, such that dose differed across sources in our assay. Controlling for dose experimentally requires extraction of endospores from soil, which results in ∼60% loss of endospores in soils with minimal amounts of clay (Costa *et al*., 2006). Our field sites consisted of ∼20% clay, preventing efficient extraction of endospores (USDA Natural Resources Conservation Service., 2025). We thus did not control endospore abundance in the experiment and instead accounted for variation among parasite sources by including parasite source as a main effect in our statistical models (Liu *et al*., 2019; Mundim and Gibson, 2022; Gibson *et al*., 2024).

### Attachment assay

To test for a G_H_xG_P_xE interaction, we evaluated these five parasite sources for their ability to attach to each of the six isofemale host lines at three temperatures. Each combination of host, parasite, and temperature (90 total) was replicated up to three times in our attachment assay (Figure A1). For each replicate, we measured attachment rate - the proportion of hosts with endospores attached - and attachment load - the number of endospores attached per host with one or more endospores attached. Attachment rate was our primary focus because it reflects the potential for infection by *P. penetrans;* a low attachment rate indicates low potential for infection and limited parasite fitness. Attachment load reflects the extent of attachment given that a host is susceptible to attachment; it provides a secondary estimate of host-parasite compatibility.

We assayed attachment rate and load in a single assay in which nematodes were shaken in water with endospores to promote contact. To set up the assay, we extracted eggs from each isofemale line according to McClure, Kruk and Misaghi (1973). We washed plant roots and cut them into 1-inch pieces. We placed the root pieces into 250mL flasks, filled them with 10% bleach, and shook them for 2 minutes to dissolve the gelatinous matrix and release the eggs. We washed the roots over stacked 120 and 500 mesh sieves to wash and isolate eggs. We collected eggs from the 500-mesh sieve and poured these into 300mL beakers to hatch at 28° C over the course of ∼5 days. We suspended hatched J2s in water at the start of the experiment.

To create endospore suspensions for each parasite source, we mixed the dry soil thoroughly then combined 30cm^3^ of soil with 200 mL of water. We mixed this solution vigorously then allowed the mixture to settle for 5 seconds to remove larger particles of soil or debris, before pouring the top 100 mL into 250mL flasks. We repeated this process until we had 54 flasks of 100 mL of endospore suspension for each parasite source (3 replicates for 6 host lines at 3 temperatures).

To initiate the experiment, we added 10 mL of homogenized J2 suspension to a 250 mL flask containing 100 mL of endospore suspension. We repeated this until each isofemale line was ultimately distributed into 45 flasks (3 replicates for 5 parasite sources at 3 temperatures). We shook the flasks at 180 RPM for 48 hours in incubators set to 25°C, 30°C, or 35°C. These temperatures were chosen to reflect the conditions the host and parasite populations experience in their native field environments (Natural Resources Conservation Service, 2025). In addition, prior studies reported that mean attachment phenotypes varied over this temperature range, supporting the potential for specificity to also vary (Hatz and Dickson, 1992; Chen and Dickson, 1997; Freitas, Mitchell and Dickson, 1997). After 48 hours, hosts were isolated by centrifugal flotation and stored at 4°C for one to six days before being examined for attached endospores using an inverted microscope at 40x magnification (Timper *et al*., 2001). For each flask, we counted the number of endospores per host for up to 50 hosts to estimate attachment rate and load. We ultimately counted an average of 30.9 ± 1.2 (standard error of the mean) hosts per flask.

This experiment was done in a block structure, where each block comprised three replicate flasks for each of two host lines paired with each parasite source over the three temperatures (90 flasks per block). Due to low numbers of hosts for host line 5, we dropped 23 replicate flasks containing this line. We also dropped one replicate flask for host line 4 (Figure A1). We ultimately measured attachment rate and load for 7,620 nematodes in 247 experimental replicates.

### Statistical Analysis

Our goal was to determine whether host–parasite specificity varied across temperatures. To assess this, we estimated attachment rate and attachment load for each combination of host line and parasite source and compared these estimates across temperatures. Parasites are considered more successful in combinations with higher attachment rates and loads.

We conducted all analyses in R version 4.3.3 using the package *tidyverse* to manipulate and plot data (R Core Team, 2018; Wickham *et al*., 2019). We fit models using the packages *lme4* and *glmmTMB* (Douglas *et al*., 2015; McGillycuddy *et al*., 2025). We assessed model fits using the package *performance* (Ludecke *et al*., 2021) and tested for over- and underdispersion. We compared candidate models of varying complexity using AIC. We could not independently assess the effect of experimental block, because a given host-parasite-temperature combination was not tested independently across blocks. Accordingly including block as a fixed effect did not improve model fit, and we excluded it from our models. Because we had to drop certain combinations including host line 5, our dataset was unbalanced, and our analyses had inflated degrees of freedom. We therefore repeated our analyses excluding host line 5. Our results were qualitatively similar, so we report the results of analyses with all host lines included in the main text, while results of the reduced analyses are included in the Appendix.

We first evaluated variation in attachment rate. Attachment rate is the proportion of hosts with at least one attached endospore, out of the total hosts counted in a sample. We fit a generalized linear mixed model with a binomial error distribution to the number of nematodes with endospores out of the total examined per flask (i.e., a grouped binomial). This approach accounts for differences in sample size among flasks (Cronin *et al*., 2010). We included a unique ID for replicate flask as an observation-level random effect to correct for overdispersion (Bolker, 2015). The full model included fixed effects for host line, parasite source, and temperature as factors and all two- and three-way interactions among these factors. We also analyzed attachment rate separately at each temperature to test for the presence of host-parasite specificity. These three models included host line, parasite source, and their interaction as fixed effects, plus a random effect for replicate flask to correct for overdispersion. In all models, the fixed effect of parasite source controlled for variation among parasite sources that was independent of host line, including differences in both intrinsic infectivity and endospore dose. After accounting for these intrinsic differences among parasite sources, terms including an interaction with parasite source allowed us to evaluate changes in the relative performance of parasite sources across temperatures (G_P_xE), host lines (i.e., G_H_xG_P_), or both (G_H_xG_P_xE).

To complement our results for attachment rate, we also evaluated variation in attachment load. Attachment load is the number of endospores attached per host, excluding hosts with no endospores attached. We analyzed attachment load by fitting a generalized linear mixed model with a negative binomial distribution truncated to exclude zeros (family = truncated_nbinom2). We detected underdispersion in this model. Because alternate distributions always led to overdispersion, increasing the likelihood of a type 1 error, we retained the negative binomial distribution. Thus, results of this analysis may be conservative. We included a unique ID for replicate flask to account for non-independence of hosts from the same flask. Fixed effects were as described for attachment rate.

Following on the results of these analyses, we quantified the variance that could be attributed to host, parasite, and interaction effects in these models. To do this, we re-fit models with each factor and interaction term as a random effect, with the exception of temperature. We retained temperature as fixed because we specifically controlled for it in our experiment. We then estimated the relative contribution of each term to the total variation accounted for by the model.

To better understand how G_H_xG_P_xE interactions influence host-parasite specificity, we conducted follow-up analyses to determine how specificity changed across temperatures. Our goal was to understand the extent to which observed G_H_xG_P_xE effects could be attributed to 1) shifts in rank order, such that host-parasite combinations that are relatively successful in one temperature are unsuccessful in another, or to 2) changes in the strength of specificity across temperatures, such that the variance attributed to the G_H_xG_P_ interaction is greater in one temperature than another. For these analyses, we report data without host line 5 to ensure a balanced design. First, we tested for shifts in the rank order of host-parasite combinations across temperatures by estimating Kendall’s rank correlation coefficient (_Ʈ_) between each pair of temperatures. An estimate of 1 indicates that rank order is conserved across temperatures, 0 indicates independence, and -1 indicates rank order is inverted (Okoye and Hosseini, 2024). Second, we compared the relative contributions to the G_H_xG_P_xE effect of rank order shifts and changes in the strength of specificity. Modifying the approach of Fry, Heinsohn and Mackay (1996), based on Robertson (1959) and Cockerham (1963), we expressed the variation in attachment rate among host-parasite combinations across temperatures as:

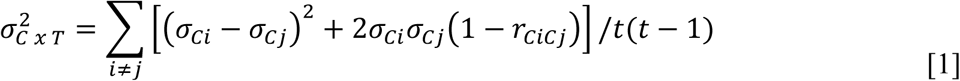

Here, *t* is the number of temperature conditions, σ_*ci*_ and σ_*cj*_ are the square-roots of the variance components among host-parasite combinations measured separately in temperatures *i* and *j*, respectively, and *r*_*cicj*_ is the variance component correlation of combinations across the two temperatures, which we calculated as:

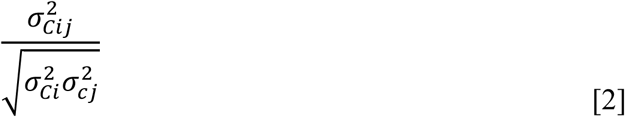

where 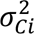 and 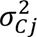 are the variance components due to host-parasite combinations measured separately in temperatures *i* and *j*, respectively, and 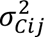 is the variance component due to host-parasite combinations in a model that combines data from temperatures *i* and *j*. This correlation approaches 1 if combination means covary strongly across temperatures. The first term in equation 1 reflects the change between temperatures in the variance attributed to host-parasite combinations; a relatively large value indicates that the strength of specificity differs substantially between temperatures. The second term reflects consistency of relative combination performance across temperatures; a relatively large value indicates that relative performance of combinations in one temperature is not predictive of relative performance in another temperature, consistent with shifts in rank order. This latter term reflects the potential for environmental conditions to alter the direction of selection. We applied this variance partitioning approach only to attachment rate, for which the G_H_xG_P_xE effect contributed substantial variance.

## RESULTS

To evaluate the effect of the environment on specificity in the interaction of *Pasteuria penetrans* with *Meloidogyne arenaria,* we measured the rate and load of *P. penetrans* endospore attachment for six lines of *M. arenaria* paired with five sources of *P. penetrans* at three temperatures. We present the main effects of temperature, host line, and parasite source, followed by the effect of temperature on host-parasite specificity (G_H_xG_P_xE) for attachment rate, then attachment load.

### Attachment Rate

The best model for attachment rate included all terms, including a three-way interaction of host line, parasite source, and temperature (ΔAIC = 13.0, Table A2). The attachment rate of *P. penetrans* varied with temperature (χ² = 62.2, *df* = 6, p < 0.001, Table A3). Attachment rate was higher in the high temperature treatment (mean ± standard error of the mean: 77.2 ± 0.9%) than in the low (67.7 ± 0.9%) and medium temperature treatments (63.4 ± 0.9%, Figure 1).

**Figure 1.**
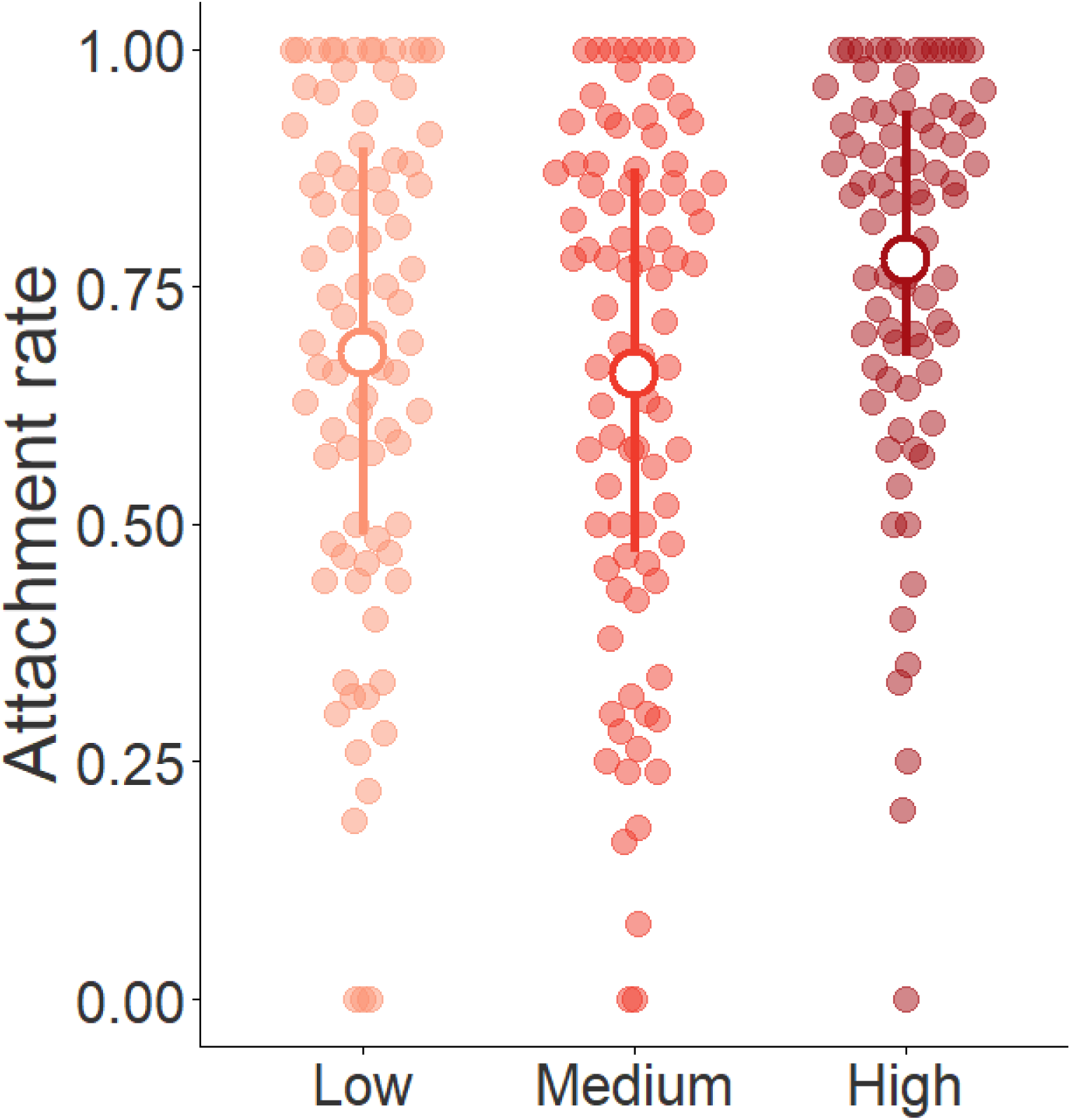
A**ttachment rate varies with temperature.** This figure shows the distribution of attachment rates, where each point represents a replicate flask at each temperature (∼90 flasks per temperature). Attachment rate is the proportion of hosts per flask with endospores attached; higher values indicate combinations with a higher probability of infection. White points indicate mean attachment rates for each temperature, and error bars indicate the interquartile range.

Attachment rate also varied with the main effects of host line (*χ²* = 96.6 *df* = 11, *p* < 0.001) and parasite source (*χ²* = 388.1, *df* = 8, *p* < 0.001) (Table A3). Across host lines, the mean attachment rate ranged from a low of 60.8 ± 1.3% (line 3) to a high of 80.6 ± 2.8% (line 5). This main effect of host line reflects intrinsic variation in host susceptibility to attachment, independent of the parasite source. Across parasite sources, the mean attachment rate ranged from a low of 46.3 ± 1.2% (source 4) to a high of 84.9 ± 0.9% (source 2). This main effect of parasite source could arise from intrinsic variation in infectivity, independent of host line, as well as variation in endospore dose.

We detected a three-way interaction of host line, parasite source, and temperature (χ² = 99.1, *df* = 36, p < 0.001) consistent with a G_H_xG_P_xE interaction for attachment rate (Figure 2, A2, Table A2, A3). This three-way interaction of host, parasite, and temperature contributed 11.1% of the variance in attachment rate accounted for by our model factors. In this full model, including data from all three temperatures, the two-way interaction of host line and parasite source (i.e., G_H_xG_P_) did not account for any variance in attachment rate (Table A4).

**Figure 2.**
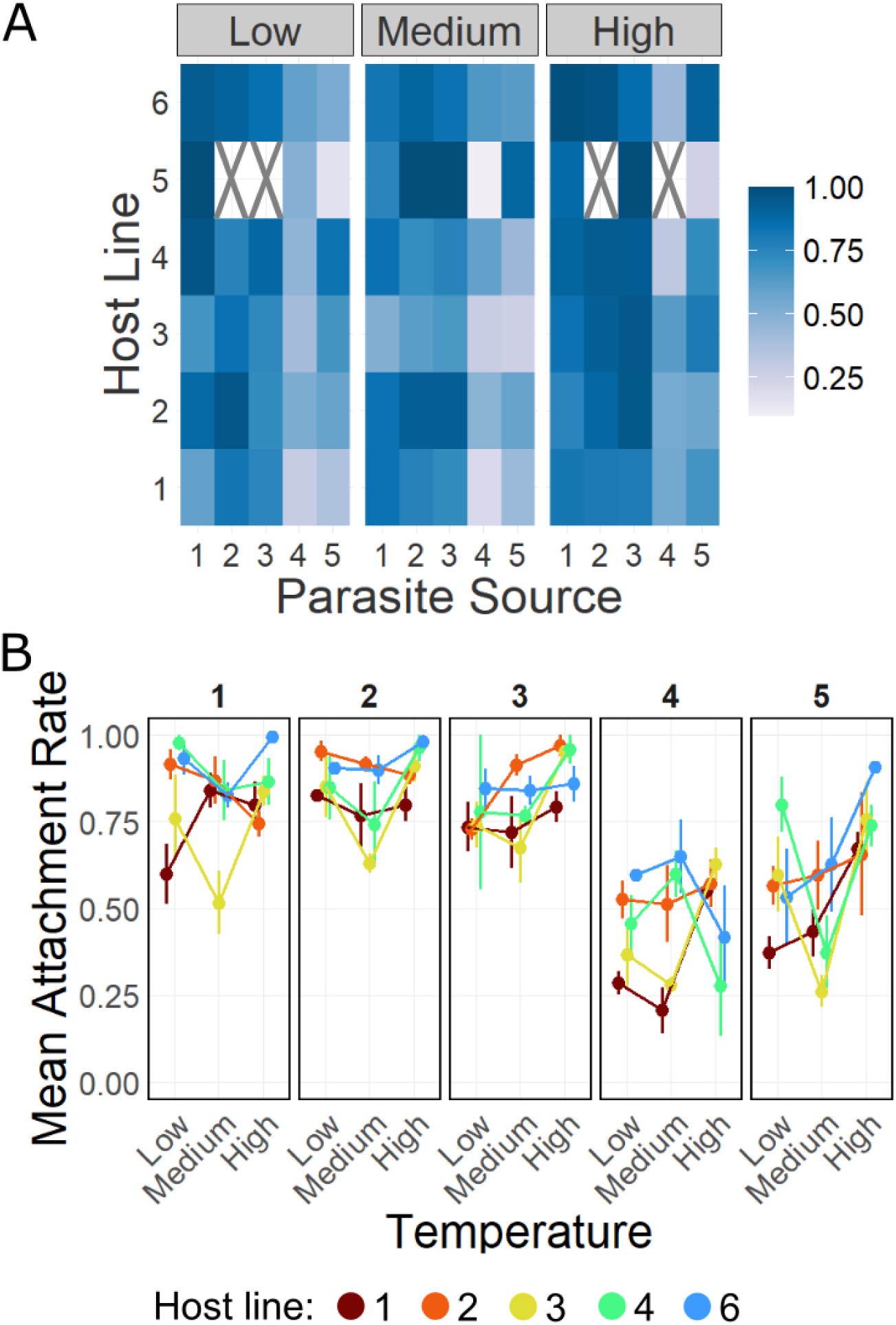
Attachment rate specificity varies with temperature. **A.** Squares represent the mean attachment rate for a single host-parasite combination at a given temperature. Host lines are ordered by row, parasite sources are ordered by column, and temperatures are split by panel. Darker squares indicate higher attachment rates. Squares with an X represent combinations with no data. **B.** Panel B shows attachment rate broken down by parasite source. It contains the same data as panel A, with the exception of host line 5, which is excluded due to having no or few replicates for many combinations. Each panel shows results for a single parasite source, and each point indicates the mean attachment rate with standard error for a host line at low, medium, and high temperatures. See Figure A2 in the Appendix for all host lines by parasite and block. Statistical results are consistent with and without host line 5.

To understand how attachment specificity changed between temperatures, we evaluated the strength of host-parasite specificity (G_H_xG_P_) at each temperature (Table A5). Attachment rate varied with the interaction of parasite source and host line at high (χ² = 51.5, *df* = 18, p < 0.001) and low temperatures (χ² = 43.1, *df* = 18, p < 0.001), but not at medium temperature (χ² = 30.5, *df* = 20, p = 0.06) (Figure 2). These results suggest that the G_H_xG_P_xE interaction detected in the full model arose in part from variation in the strength of specificity across temperatures.

To determine if temperature caused shifts in the relative performance of host-parasite combinations, we estimated the correlation across temperatures of the ranks of host-parasite combinations for attachment rate (Figure 2B). Rank correlation coefficients were moderately positive for all pair-wise combinations, indicating that the rank orders of host-parasite combinations were partially conserved between temperatures (Kendall’s _Ʈ_ = 0.56 for the correlation of low to medium temperature; 0.41 for medium to high; 0.46 for low to high, p ≤ 0.004 across comparisons).

To compare the relative importance of changes in rank order to changes in specificity for the observed G_H_xG_P_xE effect, we partitioned the variance component attributed to the three-way interaction according to equations 1 and 2. We found that changes among temperatures in the strength of host-parasite specificity (G_H_xG_P_) accounted for 42% of the variance, while 58% of the variance was attributed to the lack of correlation in the performance of host-parasite combinations across temperatures.

### Attachment Load

We then examined variation in attachment load, or the number of *P. penetrans* endospores attached to susceptible hosts. The best model for attachment load included all terms, though support for the three-way interaction was weak (ΔAIC = 2.6, Table A6). Attachment load varied with temperature (χ² = 24.2, *df* = 2, p < 0.001, Table A7). These effects were slight: median load was 10 endospores per host at the high temperature, slightly higher than the median load of 7 endospores at low and medium temperatures (Figure 3).

**Figure 3.**
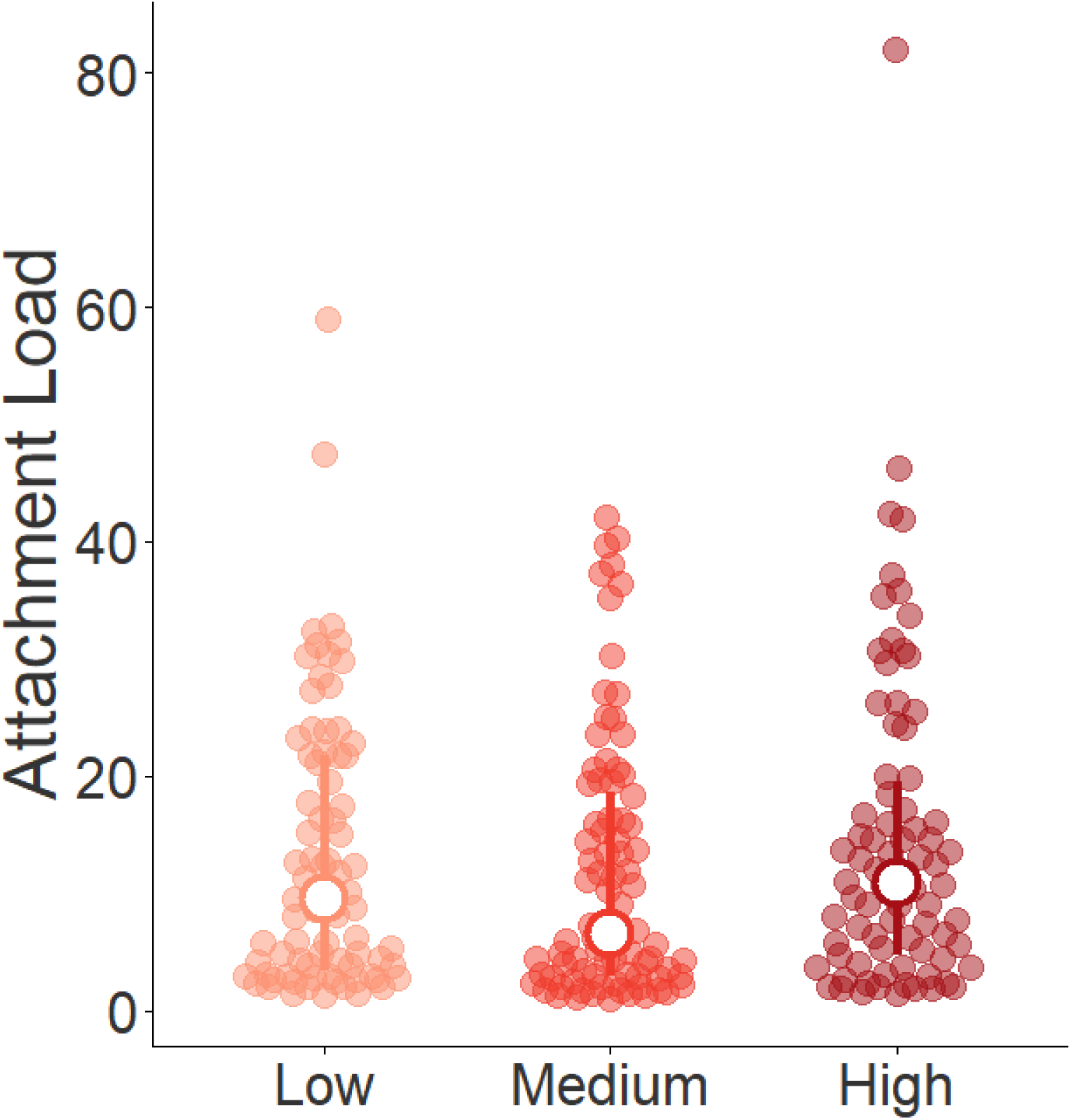
Attachment load varies with temperature. This figure shows the distribution of mean attachment load for each replicate flask at each temperature (∼90 per temperature). Attachment load is the number of endospores attached per host with one or more endospores attached. White points indicate median load across flasks for each temperature, and error bars show the interquartile range.

Attachment load also varied with the main effects of host line (χ² = 103.1, *df* = 5, p < 0.001) and parasite source (χ² = 926.7, *df* = 4, p < 0.001)(Table A7). Across host lines, median attachment load ranged from 3 (line 3) to 14 endospores per host (line 6). Across parasite sources, median attachment load ranged from 2 (source 4) to 22 endospores per host (source 3).

We detected a three-way interaction of host line, parasite source, and temperature (χ² = 84.8, *df* = 36, p < 0.001) (Figure 4, Table A6, A7). This effect was not as strong as that observed for attachment rate, with the three-way interaction of host, parasite, and temperature accounting for 6.2% of the variance in attachment load accounted for by our model factors (Table A8).

**Figure 4.**
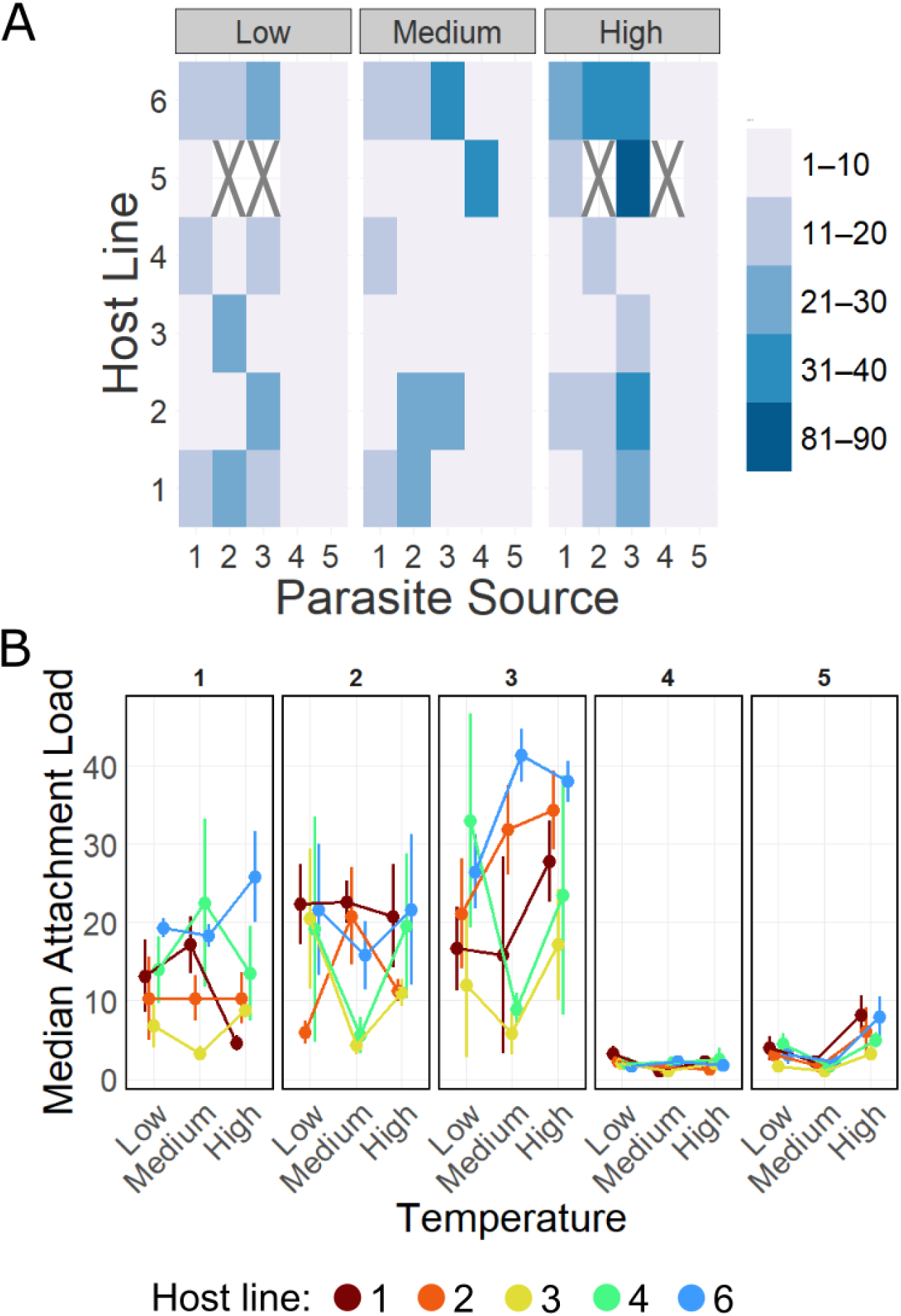
Attachment load specificity varies with temperature. **A**. Squares represent the median attachment load for a single host-parasite combination at a given temperature. Host lines are ordered by row, parasite sources are ordered by column, and temperatures are split by panel. Data is binned in intervals of ten, so each color represents a range of endospore counts. Squares with an X represent combinations with no data. **B.** Panel B shows attachment load broken down by parasite source. It contains the same data as panel A, with the exception of Host line 5, which is excluded due to having no or few replicates for many combinations. Each panel shows results for a single parasite source, and each point indicates the average of median endospore counts per flask plus standard error for a host line at low, medium, and high temperatures. Hosts with no endospores attached are excluded. See Figure A3 in the Appendix for all host lines by parasite and block. Statistical results are consistent with and without host line 5.

As we did for attachment rate, we conducted follow-up analyses to understand how attachment load specificity changed between temperatures. We first examined the strength of host-parasite specificity (G_H_xG_P_) at each temperature (Table A9). Attachment load varied with the interaction of parasite source and host line at low (χ² = 37.9, *df* = 18, p = 0.004) and medium temperatures (χ² = 68.0, *df* = 20, p < 0.001). We did not detect an interaction of parasite source and host line at high temperature, after Bonferroni correction (χ² = 30.8, *df* = 18, p = 0.030; adjusted α=0.017 for three tests). Second, we evaluated shifts in rank order among host-parasite combinations (Figure 4B). As for attachment rate, rank correlation coefficients for attachment load were moderately positive, again indicating some conservation of rank order between temperatures (Kendall’s _Ʈ_ = 0.64 for the correlation of low to medium temperature; 0.69 for medium to high; 0.67 for low to high, p<0.001).

## DISCUSSION

A proposed challenge to the Red Queen Hypothesis is that the environment may change the specificity of selection between hosts and parasites (Thomas and Blanford, 2003; Lazzaro and Little, 2008; Laine, 2009; Wolinska and King, 2009; Mostowy and Engelstädter, 2011). There is, however, limited evidence that the environment can cause such changes in specificity. Our study provides initial evidence that environmental factors, here temperature, may indeed alter specificity in the interaction of *Meloidogyne arenaria* and *Pasteuria penetrans*. We suggest that follow-up studies using controlled doses of *P. penetrans* isolates or clones are needed to fully understand the magnitude of G_H_xG_P_xE interactions in this system.

We first evaluated the main effect of temperature on parasite success. We found that the rate at which *P. penetrans* endospores attached to hosts was sensitive to temperature (Figure 1). Attachment rate was highest at 35°C, the highest temperature in our experiment. Attachment load was also highest at 35°C, though the differences between temperatures were quite small (Figure 3). Freitas, Mitchell and Dickson (1997) and Hatz and Dickson (1992) also found that attachment tended to increase with temperature, with the highest estimates at 30-35°C. Freitas, Mitchell and Dickson (1997) also tested higher temperatures and found that attachment dropped off steeply above 35°C. This finding in part motivated our choice of 35°C as the maximum temperature in our experiment, as higher temperatures would be expected to generate a G_H_xG_P_xE effect simply through a collapse in overall attachment. Importantly, mean attachment rates in our experiment remained high, over 50%, across all temperatures. Though our experiment specifically focused on the attachment process, other studies have shown that temperature also impacts *P. penetrans* within-host growth. Development is faster and proliferation is greater at 30-35°C, consistent with relatively high temperatures being optimal for *P. penetrans* (Hatz and Dickson, 1992; Chen and Dickson, 1997; Serracin *et al*., 1997).

Given this main effect of temperature, we then tested our hypothesis by evaluating the effect of temperature on attachment specificity. Prior work established that attachment of *P. penetrans* to *M. arenaria* is highly specific, consistent with a G_H_xG_P_ interaction (Stirling, 1985; Channer and Gowen, 1992; Mundim and Gibson, 2022). By measuring attachment at multiple temperatures, we found that this specificity changed with temperature (Figure 2,4,A2,A3), notably when measured as attachment rate. In analyses including all temperatures, attachment rate and load did not vary with the two-way interaction of host line and parasite source (G_H_xG_P_) (Tables A4, A8). However, the interaction of host line and parasite source contributed substantially to variation in attachment rate in analyses of individual temperatures (Tables A5, A9). These contrasting results indicate that patterns of specificity at a single temperature were not maintained across temperatures. This finding provides potential support for the idea that host-parasite specificity is sensitive to environmental context and aligns with findings in several other systems (Tétard-Jones *et al*., 2007; Bryner and Rigling, 2011; Sadd, 2011; Wendling, Fabritzek and Wegner, 2017). While G_H_xG_P_xE interactions are predicted to disrupt coevolutionary dynamics, the actual consequences of our findings for the coevolutionary dynamics of *M. arenaria* and *P. penetrans* depend on several factors. These factors include the velocity of environmental change in natural populations and the environmental sensitivity of specificity for individual genotypes of *P. penetrans* (Wolinska and King, 2009; Mostowy and Engelstädter, 2011).

We do not know the mechanisms behind the G_H_xG_P_xE interactions detected in this study. One hypothesis is that temperature alters the adhesion proteins on the coat of *P. penetrans* endospores. These proteins are thought to be critical to the attachment of endospores to the cuticle of J2 hosts (Davies *et al*., 2023), and this attachment step is where we see clear evidence for strong specificity for attachment, in both *P. penetrans* (Stirling, 1985; Channer and Gowen, 1992; Mundim and Gibson, 2022) and the related *P. ramosa* (Carius, Little and Ebert, 2001; Duneau *et al*., 2011; Luijckx *et al*., 2011, 2013). In addition, because we used parasite sources that potentially contained multiple isolates, isolates within a source could have varied in their performance across temperatures. While we expect this effect would buffer host-parasite combinations from change in attachment phenotypes across temperature, thereby diminishing our ability to detect a G_H_xG_P_xE interaction, we cannot rule out the possibility that diversity within parasite sources contributed to our results.

How a G_H_xG_P_xE interaction is generated determines the nature of the challenge it presents to the Red Queen Hypothesis. A G_H_xG_P_xE interaction can be generated in two ways. First, environmental factors may change the strength of specificity, such that a G_H_xG_P_ interaction is weaker, or absent, in one condition vs. another. Second, environmental factors may change the rank order of performance of host-parasite combinations, such that the combination expected to yield the highest infection varies across environments. The latter scenario arguably poses a greater challenge than do changes in the strength of specificity, as changes in rank order indicate that the environment alters the direction of coevolutionary selection (Wade, 2007). We evaluated both possibilities in our data, focusing specifically on attachment rate, for which the G_H_xG_P_xE effect was more substantive. First, we found evidence that the strength of specificity may have changed across temperatures. Analyzing each temperature separately, we saw that at low and high temperatures, attachment rate varied significantly with the interaction of host line and parasite source, consistent with specificity. At the medium temperature, however, this interaction was marginally significant and contributed very little to variation in attachment rate, indicating weaker support for G_H_xG_P_ specificity at this temperature (Table A5). Second, we also found evidence for shifts in the rank order of host-parasite combinations (Figure 2B, A2). For example, parasite source 4 had relatively low mean attachment rates on host lines 1 and 3 at low temperature, but relatively high attachment rates at high temperature (i.e., crossing reaction norms). Similarly, host line 6 is the most susceptible to attachment by parasite source 5 at high temperature, but relatively less susceptible to it at low temperature. For the remaining parasite sources, shifts in rank order are less evident, and mean attachment rates exceeded 50% for all host lines and temperatures. Consistent with these observations, rank correlation coefficients were positive but substantially less than 1, indicating some changes in the rank order of combinations across temperatures. Finally, by partitioning the variance associated with the G_H_xG_P_xE effect (Fry, Heinsohn and Mackay, 1996), we found that both changes in rank order and in specificity contributed to the variation among host-parasite combinations across temperatures, with changes in rank order contributing slightly more. We conclude that both changes in the strength and ranking of specificity drove the G_H_xG_P_xE effect in our data. The partial conservation of rank order in this system may not fully disrupt the expected coevolutionary dynamics. Further evaluation of its potential effect requires tailored theoretical treatment.

In other studies reporting G_H_xG_P_xE interactions, there is evidence for both changes in the strength of specificity and in rank order. The significant G_H_xG_P_xE effect in Bryner and Rigling (2011) likely arose from changes in rank order. They examined changes in the interaction of the fungus *Cryphonectria parasitica* with its hypervirus at four relevant temperatures. For growth and sporulation of fungus exposed to virus, they detected substantial crossing of reaction norms between temperatures, as well as G_H_xG_P_ interactions at each individual temperature, indicating strong specificity was maintained. The G_H_xG_P_xE effects in Tétard-Jones *et al*. (2007) and Sadd (2011) appear to reflect both changes in rank order and in the strength of specificity. Regardless of their origins, the growing evidence for G_H_xG_P_xE interactions is important, as such effects may modify coevolutionary dynamics predicted under the Red Queen. Moving forward, it is especially important for studies to quantify the relative contribution of shifts in rank order, as these are expected to have the biggest impact on coevolutionary dynamics (Mostowy and Engelstädter, 2011). Several other studies found no support for G_H_xG_P_xE effects (Cayetano and Vorburger, 2013; Gimmi and Vorburger, 2021). Notably, our findings contrast with a clear lack of evidence for G_H_xG_P_xE interactions in a closely-related system, the bacterial parasite *Pasteuria ramosa* and its host, the cladoceran *Daphnia magna* (Vale and Little, 2009; Duneau *et al*., 2011). Strong genetic specificity has been documented for this system. By crossing four isolates of *P. ramosa* with four clones of *D. magna* at three temperatures, Vale and Little (2009) found no evidence that specificity for infection changed with temperature. Similarly, Duneau *et al*. (2011) crossed two clones of *P. ramosa* with two clones of *D. magna* and found that specificity for attachment rate did not change with temperature (n=4 conditions), food (n=2), host density (n=2), or host sex (n=2). There are key differences between their studies and ours that may explain this discrepancy. Most notably, both Vale and Little (2009) and Duneau *et al*. (2011) included in their assays host-parasite combinations that were incompatible, meaning attachment or infection rate was 0%. Our combinations tended to show intermediate attachment rates, ranging from a low of 10.0% to a high of 100.0%, with a mean of 70.7 ± 2.4%. We may not have had incompatible combinations because we used parasite sources that may have contained multiple genotypes of *P. penetrans*, rather than single isolates. We previously demonstrated that increasing the genetic diversity of *P. penetrans* inocula increases attachment rates and reduces the probability that a host line has no attached endospores (Mundim and Gibson, 2022).

Consistent with this idea, the *P. ramosa* studies used either parasite clones (Duneau *et al*., 2011) or isolates (Vale and Little, 2009), which are expected to be less genetically diverse than our sources. Given this difference, our findings are not in direct conflict with Vale and Little (2009) and Duneau *et al*. (2011). Their results indicate that environmental factors cannot make an incompatible combination compatible; a qualitatively resistant host will remain resistant. Our study indicates that environmental factors can cause quantitative shifts in attachment phenotypes among compatible interactions. Together, these findings suggest that environmental factors may modulate the strength of selection but would weakly affect its direction. By this argument, we would not expect to detect a G_H_xG_P_xE interaction if we repeated our study using single isolates or clones of *P. penetrans* and included incompatible combinations.

In conclusion, this study adds to a small body of evidence showing that environmental context may alter genetic specificity, potentially influencing the maintenance of genetic diversity under negative frequency-dependent selection. We have focused specifically on G_H_xG_P_xE interactions, because significant weight has been given to these as a challenge to the Red Queen Hypothesis (Wolinska and King, 2009). We note, however, that disruptions to specificity are not the only potential impact of environmental fluctuations. Most fundamentally, the Red Queen Hypothesis and coevolutionary dynamics hinge on the strength of parasite-mediated selection (May and Anderson, 1983; Howard and Lively, 1994; Otto and Nuismer, 2004), which can be acutely sensitive to environmental context. Parasite performance varies with environmental conditions, as shown here and in many other studies (Fels and Kaltz, 2006; Nowakowski *et al*., 2016; Chen *et al*., 2024). Likewise, parasite virulence varies with environmental conditions (Ferguson and Read, 2002; Mitchell *et al*., 2005; Mursinoff and Tack, 2017). It is not clear what effect environmental variation in the strength of parasite-mediated selection has: it has been hypothesized to disrupt (Blanford *et al*., 2003), enhance (Gibson, Stoy and Lively, 2018), or have relatively little effect on coevolutionary dynamics (Mostowy and Engelstädter, 2011). These main effects of fluctuating environments are expected to occur concurrently with their interactive effects, including the G_H_xG_P_xE interactions evaluated here. It is accordingly quite difficult to make concrete predictions for the robustness of coevolutionary selection in naturally fluctuating environments (Wolinska and King, 2009). This complex reality calls for caution in drawing coevolutionary inferences from individual estimates of the environmental sensitivity of host and pathogen traits.

## ACKNOWLEDGEMENTS

We thank Patty Timper for guidance and support in field sampling, as well as Dillen Rogers for assistance with field work. We also want to thank Dave Carr for support in statistical analyses and the Gibson lab for comments on the work. We thank the numerous undergraduate students who helped with the upkeep of our host lines.

## ETHICS STATEMENT

We followed protocols reviewed and approved by the United States Department of Agriculture to prevent spread of *Meloidogyne arenaria* (USDA permit #526-23-159-97135). This work did not require approval from an animal welfare committee.

## AI STATEMENT

AI was not used in the production of this manuscript.

## DATA ACCESSIBILITY

Data and code used to produce this manuscript are available in the Dataset DOI: 10.5061/dryad.6q573n6df

## APPENDIX

**Table A1.** Host and parasite sampling locations. Host lines and parasite sources were collected from two farms in Tifton, Georgia. We selected host lines and parasite sources from different small fields (Blackshank) or from different plots within a large field (Gibbs) to increase the likelihood that samples were genetically distinct.

|  | Farm | Field or plot | Latitude, Longitude |
| --- | --- | --- | --- |
| Host Line 1 | Blackshank | Lower Field 1 | 31.502291, -83.546619 |
| Host Line 2 | Gibbs | Tubbs Field, Plot 410 | 31.433331, -83.580118 |
| Host Line 3 | Gibbs | Tubbs Field, Plot 301 | 31.433306, -83.580825 |
| Host Line 4 | Blackshank | Lower Field 4 | 31.501904, -83.546970 |
| Host Line 5 | Blackshank | Upper Field 4 | 31.500986, -83.546786 |
| Host Line 6 | Blackshank | Upper Field 4 | 31.500870, -83.547008 |
| Parasite Source 1 | Blackshank | Lower Field 3 | 31.501964, -83.546657 |
| Parasite Source 2 | Blackshank | Lower Field 5 | 31.501636, -83.546830 |
| Parasite Source 3 | Blackshank | Upper Field 3 | 31.501038, -83.546599 |
| Parasite Source 4 | Blackshank | Upper Field 5 | 31.500720, -83.546534 |
| Parasite Source 5 | Gibbs | Tubbs Field, Plot 506 | 31.433752, -83.580761 |

**Table A2.**
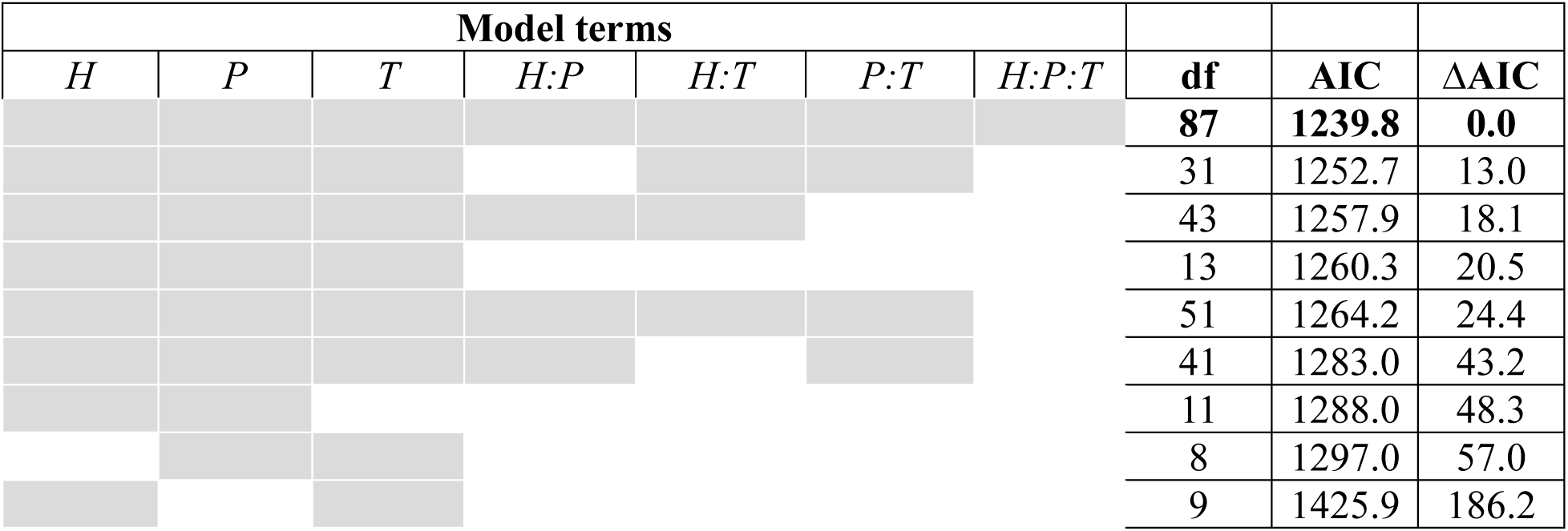
Model comparison for attachment rate. Attachment rate is the fraction of hosts with endospores attached. We compared models using AIC. All models included a random effect for replicate flask. H = host line; P = parasite source; T = temperature. Gray shading indicates a term is included in the model. The best model is bolded.

**Table A3.**
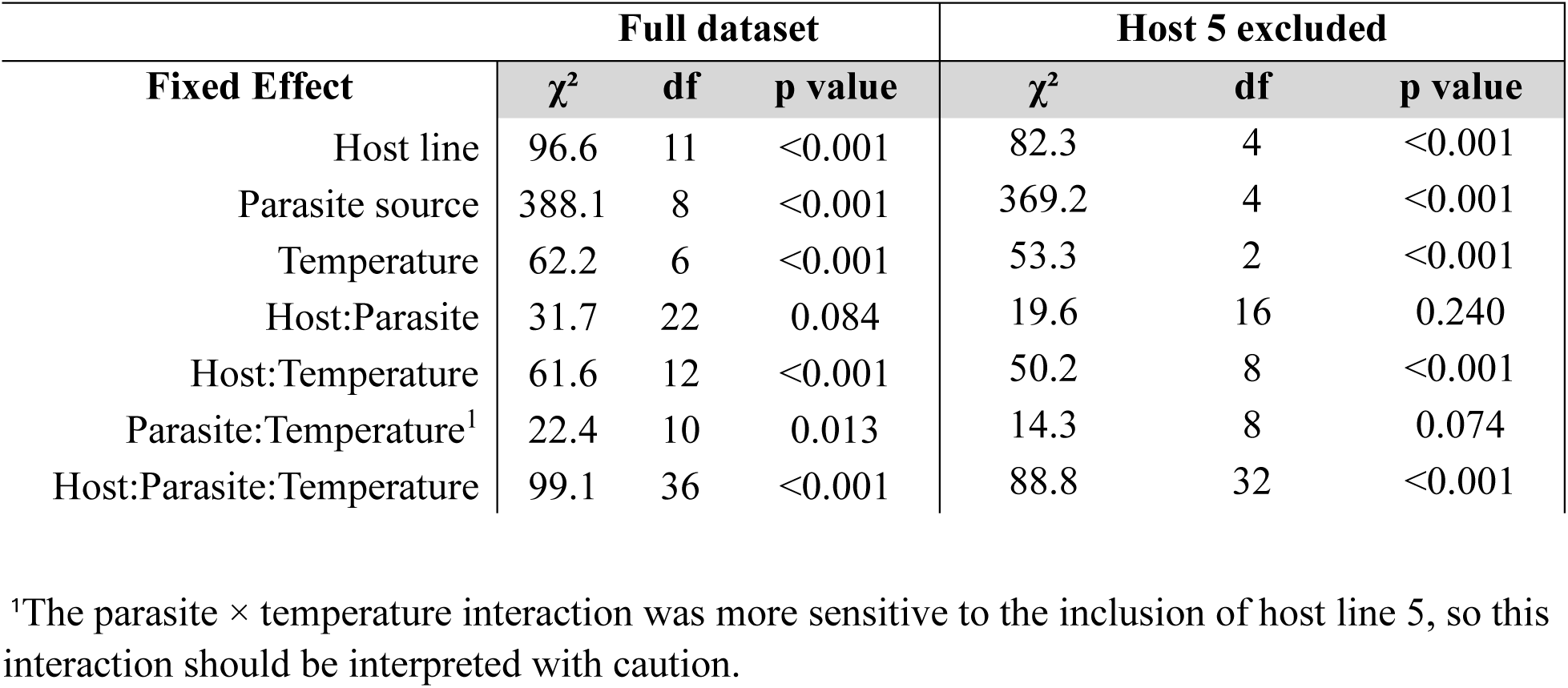
Analysis of Deviance Table for full model for attachment rate with and without host line 5. . Type II Wald χ² test. *Full model*: (Number of hosts without endospores, number of hosts with) ∼ Host line *Parasite Source*Temperature + (1|FlaskID)

**Table A4.**
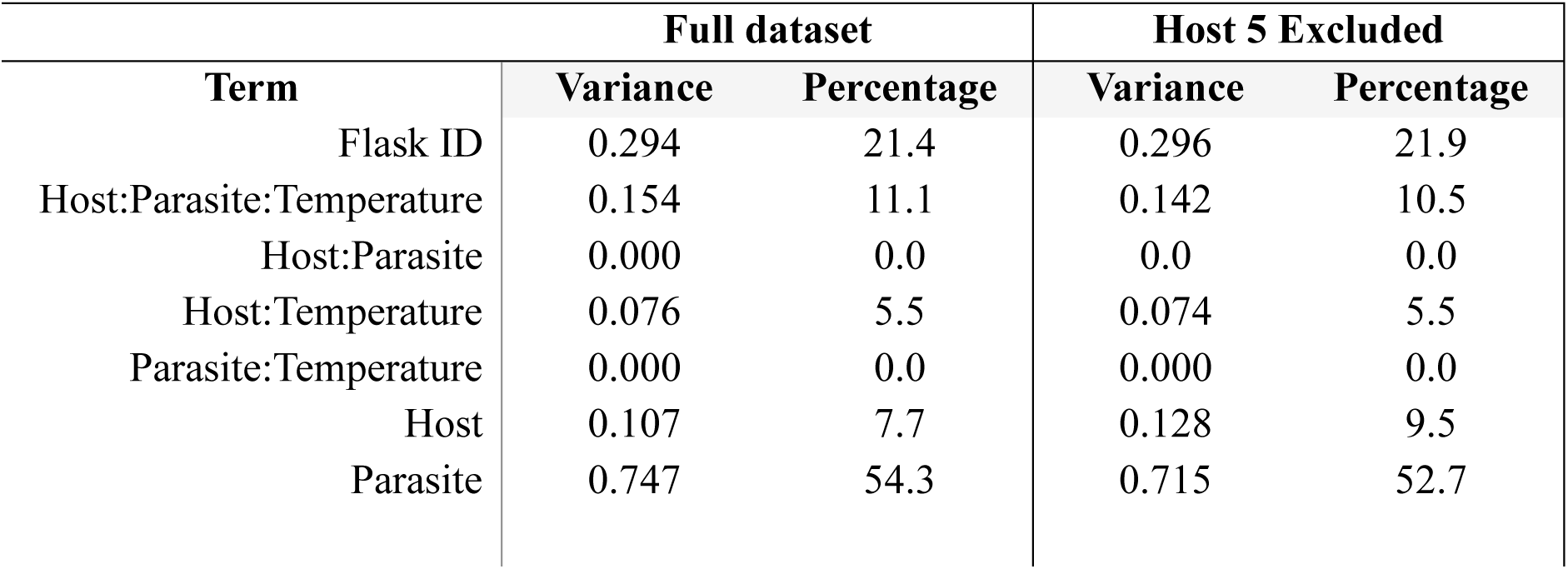
Variance components analysis for attachment rate with and without host line 5. This table shows the amount of variance attributed to each factor in the model for attachment rate.

| Term | Full dataset |  | Host 5 Excluded |  |
| --- | --- | --- | --- | --- |
|  | Variance | Percentage | Variance | Percentage |
| Flask ID | 0.294 | 21.4 | 0.296 | 21.9 |
| Host:Parasite:Temperature | 0.154 | 11.1 | 0.142 | 10.5 |
| Host:Parasite | 0.000 | 0.0 | 0.0 | 0.0 |
| Host:Temperature | 0.076 | 5.5 | 0.074 | 5.5 |
| Parasite:Temperature | 0.000 | 0.0 | 0.000 | 0.0 |
| Host | 0.107 | 7.7 | 0.128 | 9.5 |
| Parasite | 0.747 | 54.3 | 0.715 | 52.7 |

**Table A5.** Analysis of Deviance Tables for attachment rate at each temperature. . Type II Wald χ² tests. To account for multiple testing, we used a Bonferroni-adjusted α of 0.017. *Full model:* (Number of hosts without endospores, number of hosts with) ∼ Host line *Parasite Source + (1|FlaskID)

|  | *Low* |  |  |  | *Medium* |  |  |  | *High* |  |  |  |
| --- | --- | --- | --- | --- | --- | --- | --- | --- | --- | --- | --- | --- |
| | $\chi^2$ | df | p | % Variance | $\chi^2$ | df | p | % Variance | $\chi^2$ | df | p | % Variance |
| **Host Line** | 48.3 | 6 | <0.001 | 14.5 | 67.5 | 5 | <0.001 | 22.2 | 16.6 | 5 | 0.005 | 5.7 |
| **Parasite Source** | 157.1 | 5 | <0.001 | 56.0 | 127.3 | 4 | <0.001 | 49.2 | 114.4 | 4 | <.001 | 53.1 |
| **Host:Parasite** | 43.1 | 18 | <0.001 | 10.4 | 30.5 | 20 | 0.062 | 2.8 | 51.5 | 18 | <.001 | 24.6 |

**Table A6.**
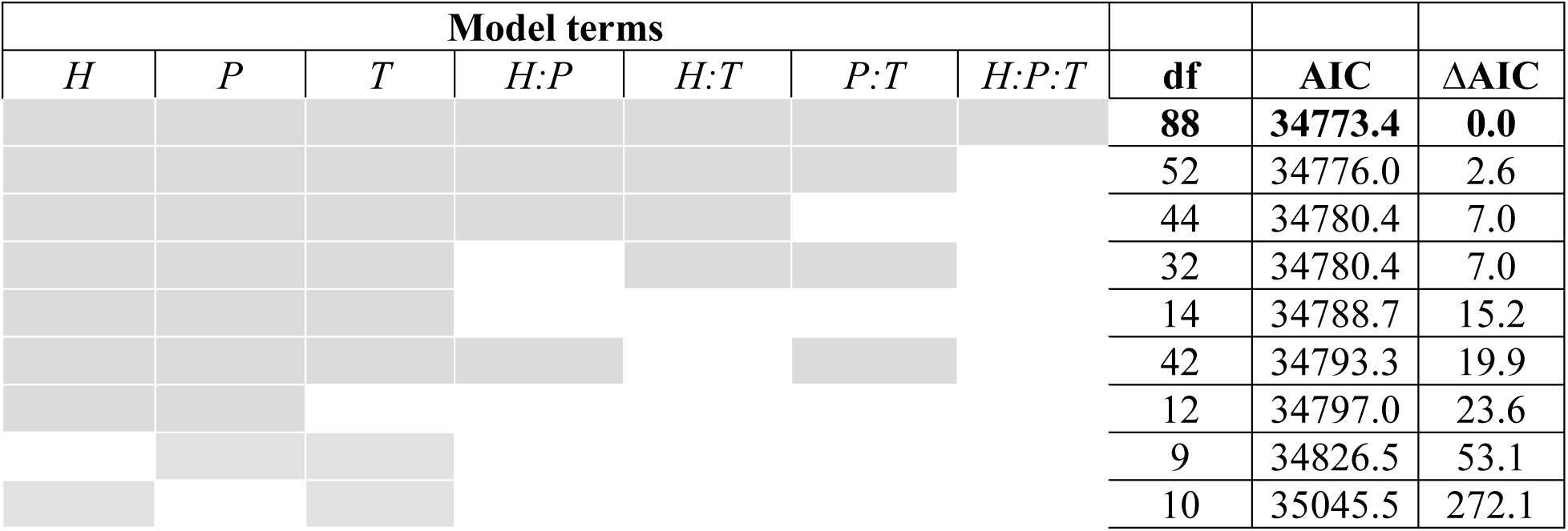
Model comparison for attachment load. Attachment load is the mean number of endospores per host, for hosts with one or more endospores attached. We compared models using AIC. All models include a random effect for replicate flask. H = host line; P = parasite source; T = temperature. Gray shading indicates a term is included in the model. The best model is bolded.

**Table A7.** Analysis of Deviance Table for full model for attachment load with and without host line 5. Type II Wald χ² test. *Full model:* Number endospores per host ∼ Host line *Parasite Source*Temperature + (1|FlaskID)

| Fixed Effect | Full dataset |  |  | Host 5 excluded |  |  |
| --- | --- | --- | --- | --- | --- | --- |
| | $\chi^2$ | df | p value | $\chi^2$ | df | p value |
| Host line | 103.1 | 5 | <0.001 | 100.0 | 4 | <0.001 |
| Parasite source | 926.7 | 4 | <0.001 | 910.0 | 4 | <0.001 |
| Temperature | 24.2 | 2 | <0.001 | 22.8 | 2 | <0.001 |
| Host:Parasite | 52.9 | 20 | <0.001 | 40.7 | 16 | <0.001 |
| Host:Temperature | 51.8 | 10 | <0.001 | 42.9 | 8 | <0.001 |
| Parasite:Temperature | 27.7 | 8 | <0.001 | 28.3 | 8 | <0.001 |
| Host:Parasite:Temperature | 84.8 | 36 | <0.001 | 74.8 | 32 | <0.001 |

**Table A8.**
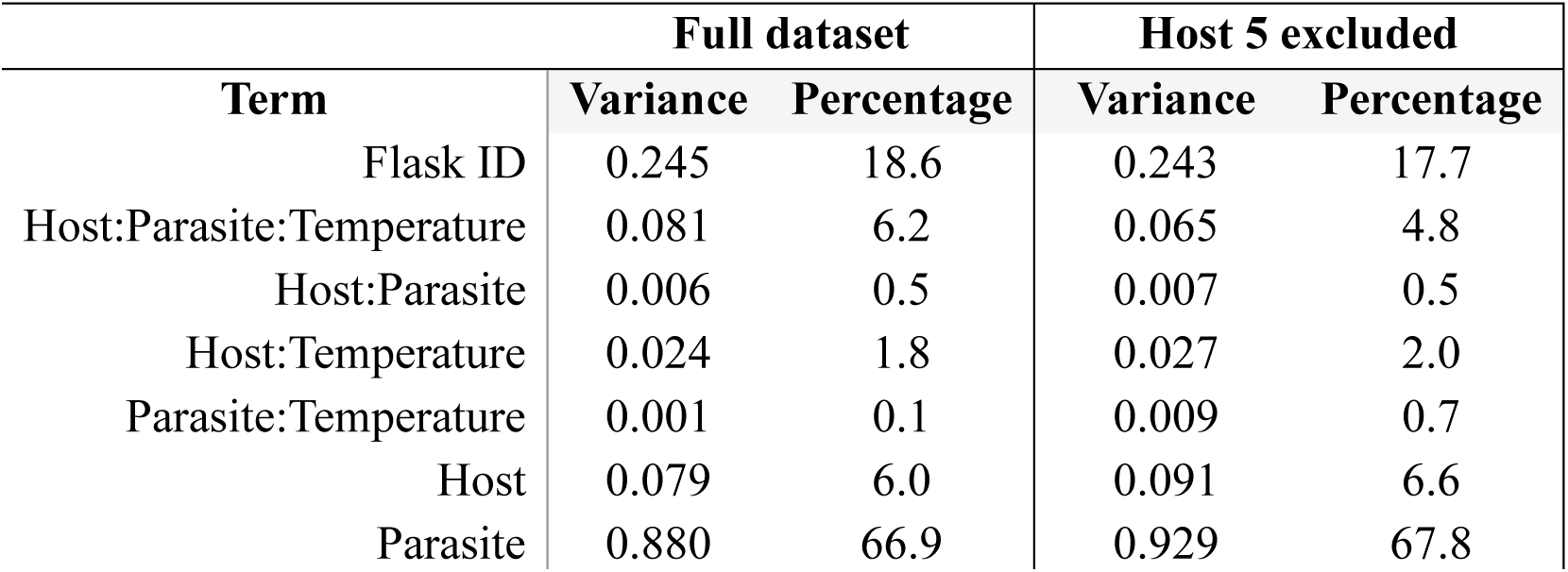
Variance components analysis for attachment load with and without host line 5. This table shows the amount of variance attributed to each factor in the model for attachment load.

| Term | Full dataset |  | Host 5 excluded |  |
| --- | --- | --- | --- | --- |
|  | Variance | Percentage | Variance | Percentage |
| Flask ID | 0.245 | 18.6 | 0.243 | 17.7 |
| Host:Parasite:Temperature | 0.081 | 6.2 | 0.065 | 4.8 |
| Host:Parasite | 0.006 | 0.5 | 0.007 | 0.5 |
| Host:Temperature | 0.024 | 1.8 | 0.027 | 2.0 |
| Parasite:Temperature | 0.001 | 0.1 | 0.009 | 0.7 |
| Host | 0.079 | 6.0 | 0.091 | 6.6 |
| Parasite | 0.880 | 66.9 | 0.929 | 67.8 |

**Table A9.**
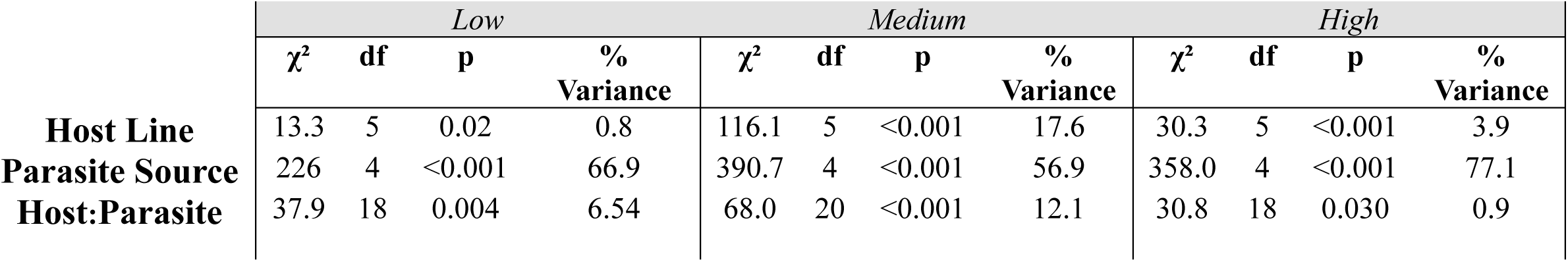
Analysis of Deviance Table for attachment load at each temperature. Type II Wald χ² tests. To account for multiple testing, we used a Bonferroni-adjusted α of 0.017. Full model: Number endospores per host ∼ Host line *Parasite Source + (1|FlaskID)

|  | *Low* |  |  |  | *Medium* |  |  |  | *High* |  |  |  |
| --- | --- | --- | --- | --- | --- | --- | --- | --- | --- | --- | --- | --- |
| | $\chi^2$ | df | p | %<br>Variance | $\chi^2$ | df | p | %<br>Variance | $\chi^2$ | df | p | %<br>Variance |
| **Host Line** | 13.3 | 5 | 0.02 | 0.8 | 116.1 | 5 | <0.001 | 17.6 | 30.3 | 5 | <0.001 | 3.9 |
| **Parasite Source** | 226 | 4 | <0.001 | 66.9 | 390.7 | 4 | <0.001 | 56.9 | 358.0 | 4 | <0.001 | 77.1 |
| **Host:Parasite** | 37.9 | 18 | 0.004 | 6.54 | 68.0 | 20 | <0.001 | 12.1 | 30.8 | 18 | 0.030 | 0.9 |

**Figure A1.**
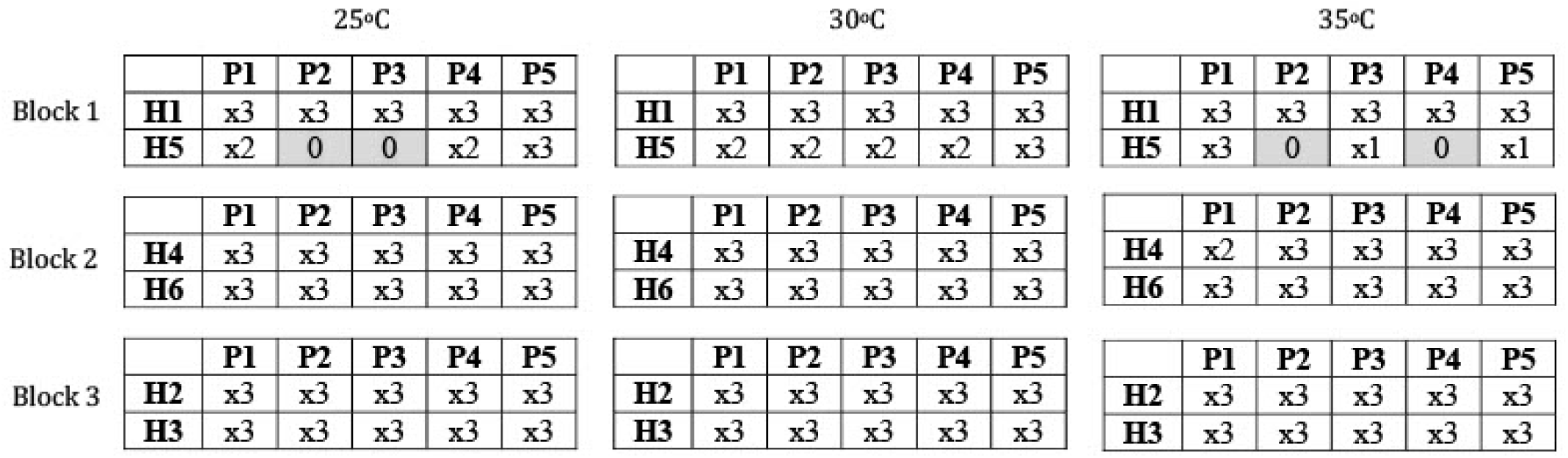
Experimental design. We selected six host lines and five parasite sources for this experiment. We paired each host line with each parasite source and replicated combinations up to three times at each of three temperatures. Cells indicate the number of replicates per combination. Host line 5 had fewer individuals, so several combinations involving this line had fewer than three replicates. The experiment was completed in three blocks, with two host lines per block.

**Figure A2.**
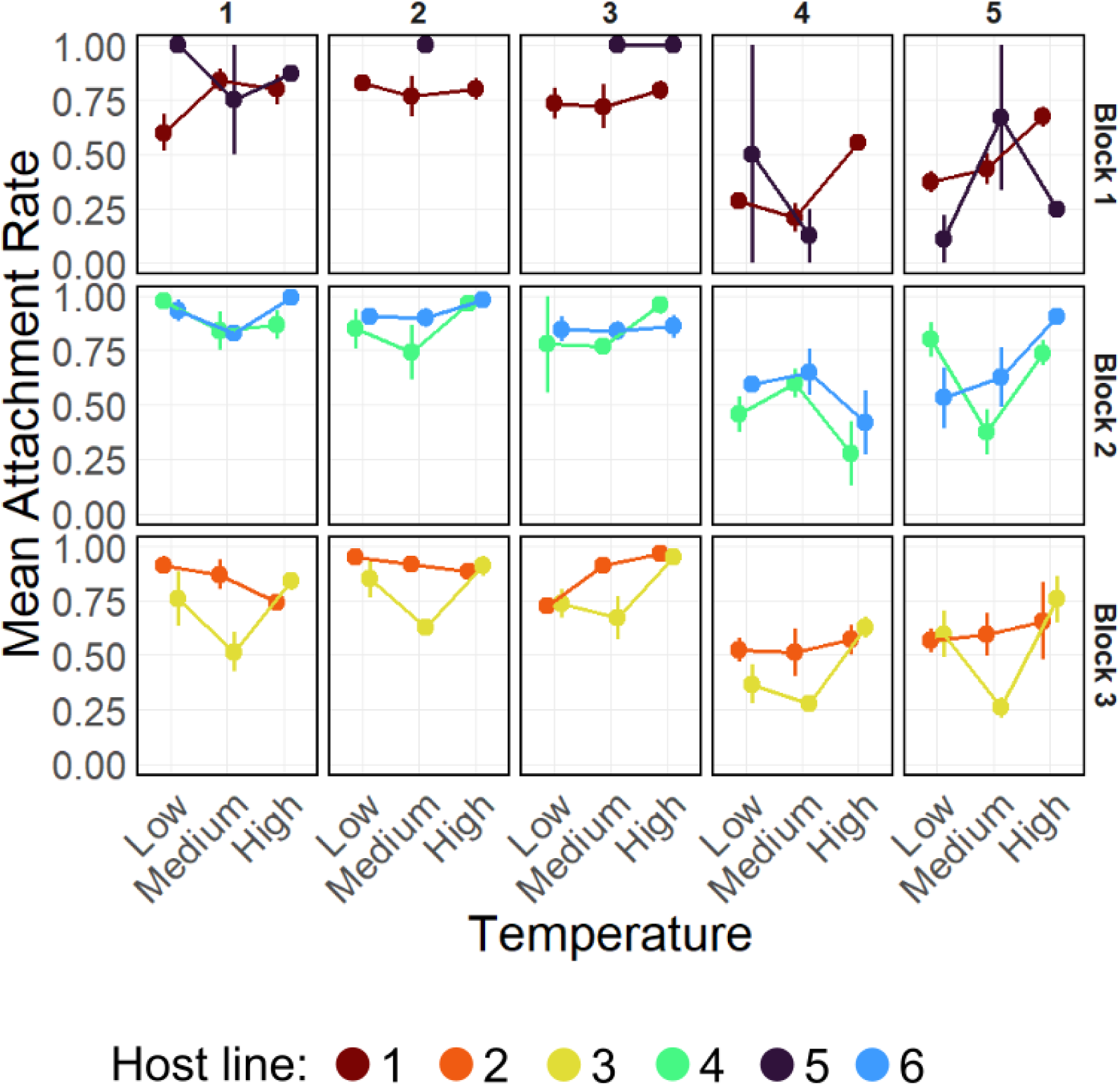
Attachment rate varies with temperature. This figure shows the mean attachment rate across low, medium, and high temperatures for each combination of parasite source and host line. This figure is identical to Figure 2b, except panels are separated by block (rows), as well as by parasite source (columns), and host line 5 is included. Points indicate means and error bars show standard error.

**Figure A3.**
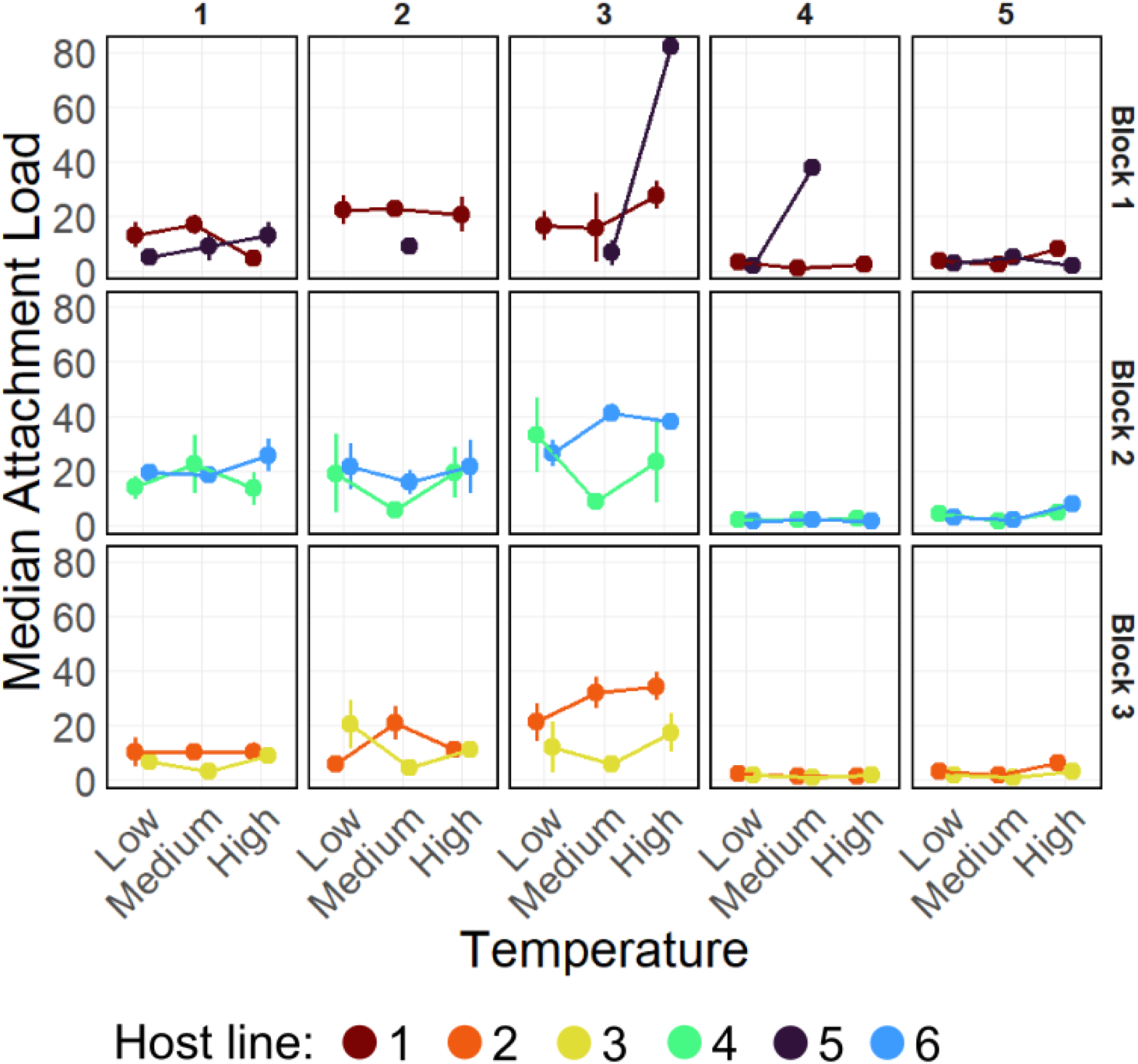
Attachment load varies with temperature. This figure shows the average of median endospore attachment load across low, medium, and high temperatures for each combination of parasite source and host line. This figure is identical to Figure 4b, except panels are separated by block (rows), as well as by parasite source (columns), and host line 5 is included. Points indicate means and error bars show standard errors.

